# Analysis of spectral and photosynthetic response variations of *Cystoseira* spp. exposed to Chromium

**DOI:** 10.1101/2025.05.20.655136

**Authors:** F. Varini, M. Penna, D. Varghese, F. P. Mancuso, M. Marcelli

## Abstract

This study explores the physiological and spectral responses of different species belonging the genus *Cystoseira* to varying concentrations of chromium (Cr) in controlled experimental conditions. The research highlights the potential of *Cystoseira* spp. as a bioindicator for monitoring heavy metal pollution in marine ecosystems. Experiments were conducted in tanks with increasing chromium concentrations (0, 0.1, 1, and 10 µmol l⁻¹), analyzing changes in reflectance, photosynthetic efficiency, and pigment composition over 14 days. Results showed dose-and time-dependent effects, with higher chromium levels causing significant photosynthetic inhibition, oxidative stress, and chlorophyll degradation. Spectral analyses identified key wavelengths (550 nm and 710 nm) correlating with chromium-induced stress, supporting their use in remote sensing for environmental monitoring. The study underscores the importance of integrating field validation and remote sensing for large-scale monitoring of marine ecosystems impacted by heavy metal contamination. These findings contribute to conservation strategies and the development of innovative tools for assessing the ecological quality of coastal waters.

## 2. Introduction

The increasing pressures exerted by human activities on marine ecosystems have highlighted the urgent need for comprehensive conservation strategies (Roberts et al., 2017). These ecosystems provide essential ecological, economic, and social benefits but are severely impacted by pollution, climate change, overexploitation of resources, and habitat destruction (Ford et al., 2022). In response, international initiatives, such as the United Nations Convention on Biological Diversity (CBD) and the European Union’s Marine Strategy Framework Directive (MSFD), emphasise the importance of monitoring and protecting marine biodiversity (Boiral and Heras-Saizarbitoria, 2017).

A key approach to assessing marine ecosystem health is using bioindicators-species or communities whose responses to environmental changes provide valuable insights into ecological conditions (Parmar et al., 2016). Among marine bioindicators, macroalgae, particularly species within the *Cystoseira* genus, have proven highly effective due to their sensitivity to environmental stressors (Pagana et al., 2024). These brown algae are foundational species in Mediterranean and Atlantic coastal ecosystems, where they form complex habitats that support a wide range of marine organisms (Mancuso et al., 2024; Lipej et al., 2023). However, their populations are in decline due to anthropogenic pressures such as pollution, habitat degradation, and climate change (Mancuso et al., 2018).

*Cystoseira* species are particularly valuable in biomonitoring programs due to their ability to bioaccumulate contaminants, including heavy metals introduced into marine environments through industrial and urban discharges (Boundir et al., 2019). Heavy metal pollution poses serious risks to marine life, disrupting physiological processes and threatening the long-term survival of sensitive species (Ali et al., 2019; Chernova and Shulkin, 2019). Studies have documented significant declines in *Cystoseira* populations in industrialized coastal areas, underscoring the urgent need for effective pollution mitigation strategies (Smith et al., 2023). Traditional monitoring methods, such as in situ surveys and laboratory analyses, provide essential data but are often labour-intensive and limited in scope (Gianni and Mangialajo, 2016). Recent advances in remote sensing technologies, including satellite imagery and hyperspectral sensing, offer new opportunities to study and map *Cystoseira* distribution across large spatial and temporal scales (Gutierres et al., 2016; Tait et al., 2019). Integrating remote sensing with ground-based validation enhances assessment accuracy, facilitating the development of targeted conservation measures (Chang et al., 2015).

The conservation of *Cystoseira* species is vital for maintaining the ecological integrity of coastal waters. These algae contribute to nutrient cycling and habitat provision, supporting marine biodiversity and local economies dependent on fisheries and tourism (Mancuso et al., 2024; Cotas et al., 2023). Protecting these habitats is both an ecological necessity and a socio-economic priority, particularly in regions where human well-being is closely linked to marine ecosystem health (Spalding et al., 2014). By combining traditional biomonitoring approaches with emerging technologies, researchers can develop more effective strategies to safeguard *Cystoseira* populations and the ecosystems they support.

## 3. The main objective of this study

To evaluate the variation of the spectral signature of *Cystoseira* spp. subjected in tanks to the exposure of the pollutant chromium at different concentrations to highlight substantial alterations for potential future applications in remote sensing.

## 4. Study Area

The coastal marine area of Civitavecchia (Lazio, Italy) is located in the northern Tyrrhenian Sea and falls in the physiographic unit M. Argentario-Capo Linaro (Fig.1). The sediments in the study area come mainly from the Mignone river and secondarily from other minor rivers, but these influence the marine sedimentation only locally and during the phases of high flood (Romão et al., 2024). The Mignone river basin (500 km^2^) is located between the Tyrrhenian margin (to the west) and the Pleistocene volcanic reliefs of the Sabatini and Cimino-Vicano mountains. Among the minor rivers that provide local contributions, the most important is the Marangone basin. Its delta is located south of the port of Civitavecchia. Although modest in size (only 23 km^2^), the Marangone stream has abandoned in its basin mines of pyrite, marcasite and galena, as well as deposits of marcasite slag and small galena pits (Kreidie et al., 2011), which strongly influence the contributions of some trace metals in the coastal area.

**Fig. 1.**
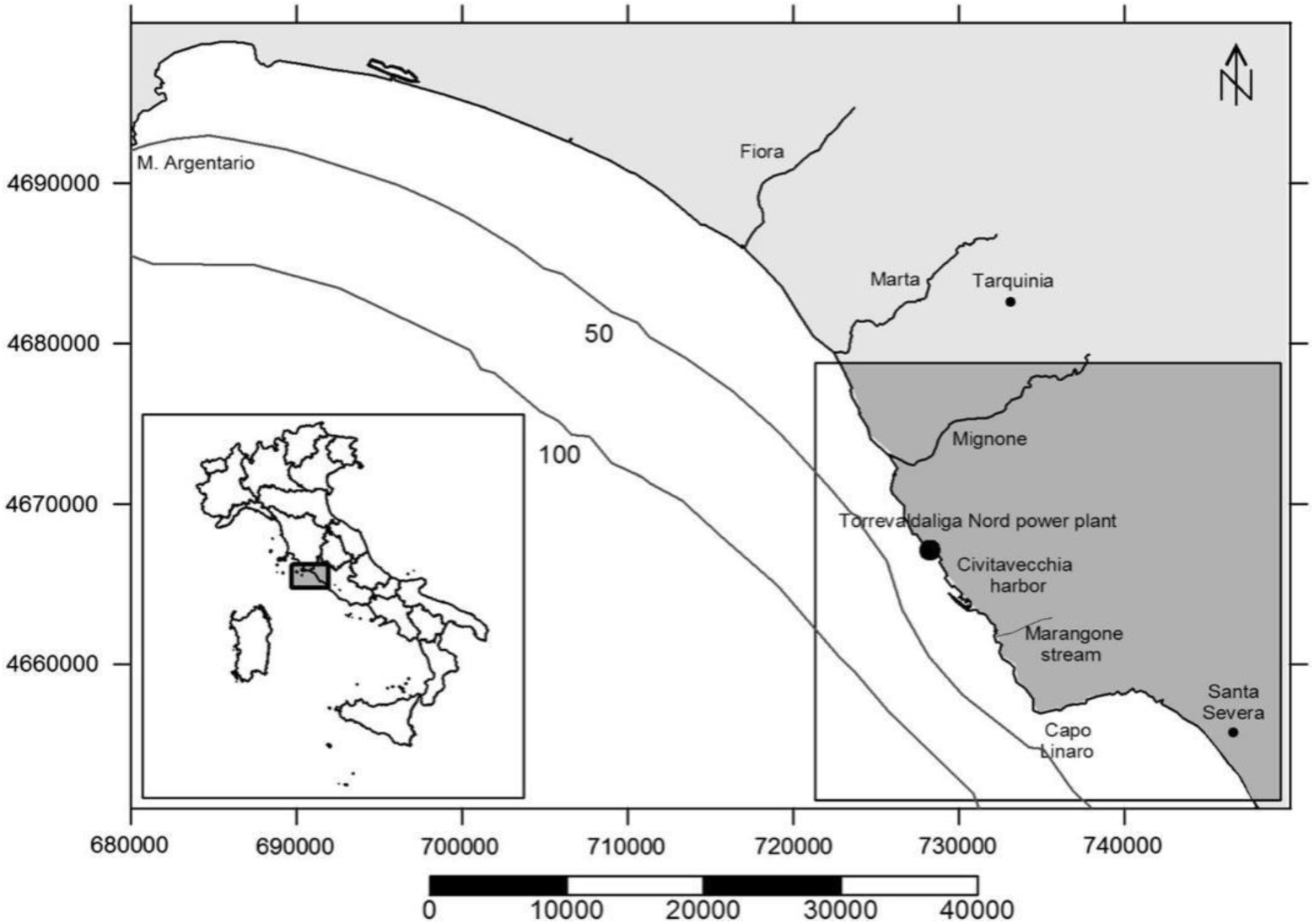
Map of Civitavecchia coast highlighting rivers, towns, industrial sites, bathymetry and study area location.

The Civitavecchia area hosts the largest energy production site in Europe, as well as one of the most important ports for cruise traffic in the Mediterranean Sea (Gobbi et al., 2020). The recent history of Civitavecchia (i.e. post-war) is marked by various industrial activities that have continuously influenced the concentrations of trace metals in all environmental compartments, particularly in the sedimentary matrix (Luciani, 2023).

The geomorphologic and anthropic characteristics of the coastal area of Civitavecchia determine a geochemical context of the environmental matrices that appears complex, given the overlap between the presence of geochemical provinces in particular of AS and Hg and the decades-long presence of industrial activities with polluting inputs of various trace metals, including those referable both to coal-fired power generation activities and others referable to maritime activities and cruise traffic. Over the years, several studies have been conducted both in the sediment matrix and in some bio-indicators in order to assess both natural and anthropogenic enrichments. Trace metal concentrations in marine sediments follow the order Mn ˃ Pb ˃ Cr ˃ Ni ˃ As ˃ Cu. The concentrations of Pb, Cr, Ni and Cu present an increasing gradient coast - wide up to the bathymetric of - 50 m, while Mn and As present maximum concentrations in two hotspots coinciding with the mouths of the Mignone and Marangone rivers, supporting the hypothesis of the main geochemical nature of these enrichments (Piazzolla et al., 2015).

The potential ecological Risk Index (RI), described by Hakanson (1980), shows a spatial distribution in which a severe toxicity hotspot is found in the area between the port of Civitavecchia and the Marangone Delta and a moderate toxicity level in the coastal area between Capo Linaro and the mouth of the Mignone river (Piazzolla et al., 2015).

A field survey was conducted to identify the presence and extent of *Cystoseira* spp. in the coastal area of Civitavecchia from 24 to 26 January 2023. In total, 4 coastal areas were investigated (fig.2).

**Fig. 2.**
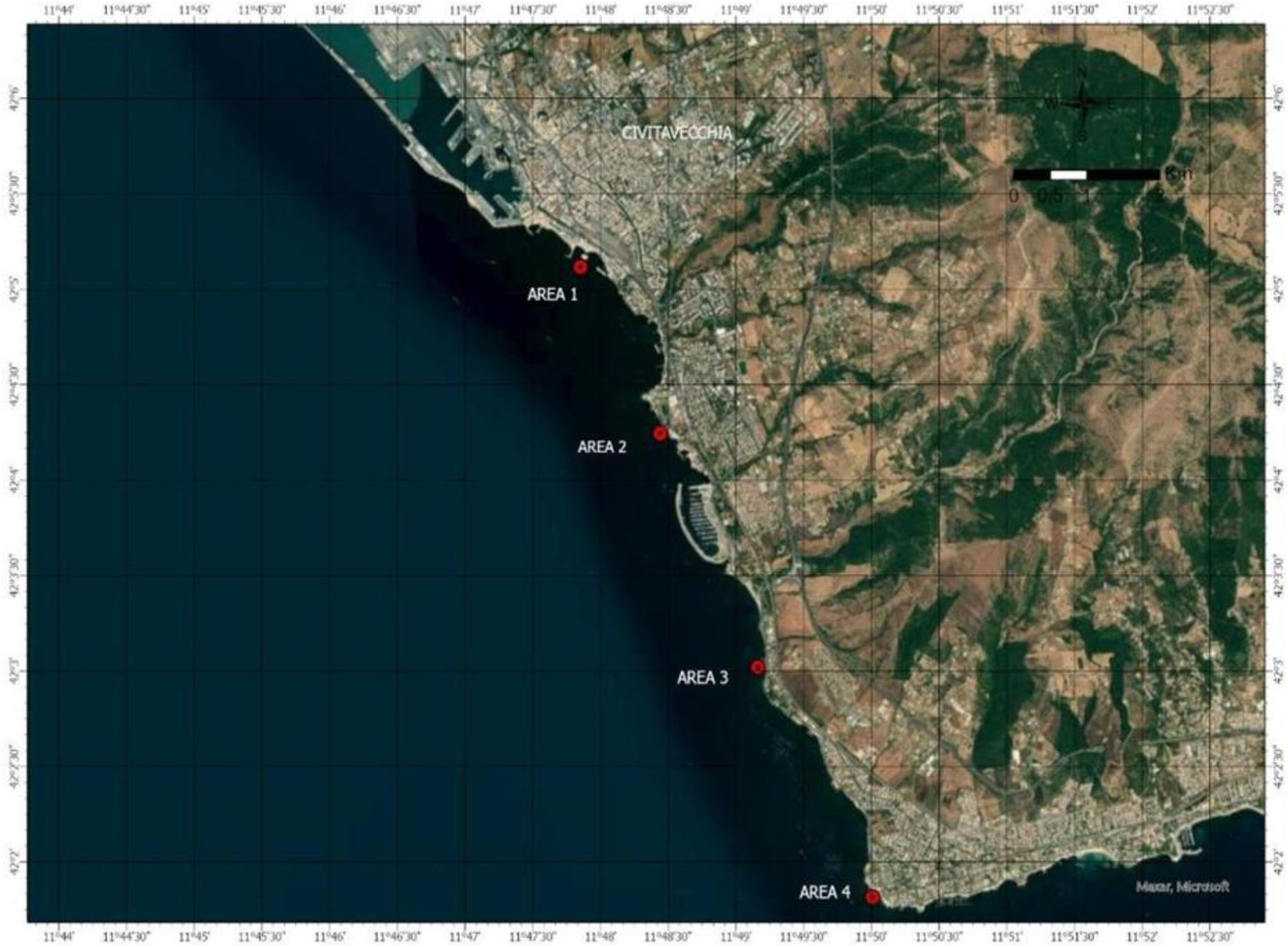
Investigated areas of *Cystoseira* spp. visual census monitoring

**Fig. 3.**
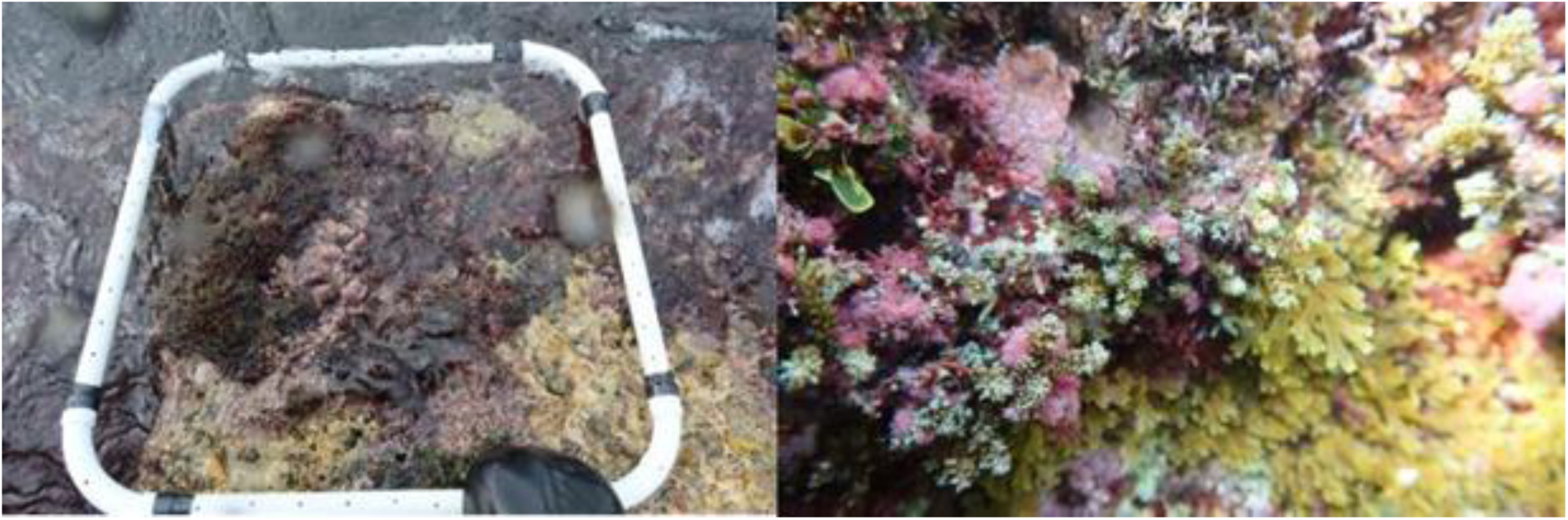
Cystoseira spp. detected within Area 3

Here we report a brief description of each area and the sampling activities carried out for the analysis of Cr in *Cystoseira* spp. samples and seawater in the area selected for the collection of samples to be exposed to the experimental enrichment assays.

Area 1

The area is located approximately 1 km southeast of the port. The coastline is characterised by hard substrates, mainly of anthropogenic nature. In this area, *Cystoseira* spp. was absent. Algal communities were mainly represented by coralline algae (*Ellisolandia elongata* (J. Ellis & Solander) K. R. Hind & G. W. Saunders, 2013*, Jania* sp. J. V. Lamouroux, 1812), brown algae *Dictyota dichotoma* (Hudson) J. V. Lamour, 1809 and red algae *Asparagopsis armata* Harvey, 1855.

Area 2

In the area located approximately 2 km southeast of the port of Civitavecchia, on the southern side (42.07061 N, 11.80758 E) *Cystoseira* species were also absent here.

Area 3

In the area located about 5 km south-east of the Port of Civitavecchia, surveys were conducted along transects parallel to the coast, inspecting the natural bedrock and the outcropping (Roman) anthropogenic structures. At the site’Molo L’ (42.05034 N, 11.81955 E) the presence of *Cystoseira* spp. was confirmed. Along the external side of the structure (about 30 m long) a band of *Cystoseira* spp. is present, more abundant. On the boulders north of the pier, *Cystoseira* spp. is present with a reduced coverage.

A hundred metres further south, at the site’Peschiera della villa romano’ (42.04882 N, 11.81942 E), isolated patches of *Cystoseira* spp. were observed.

In correspondence with the areas indicated in Fig. 4 replicates of *Cystoseira* spp. (500 g for each sample) and seawater samples (250 ml for each sample) were collected and immediately analyzed for the detection of Cr concentrations. Additional samples of *Posidonia oceanica* were collected in proximity to the *Cystoseira* spp. sampling sites for the same set of analyses. Laboratory analyses were performed according to the EPA method 7199:1996 for total Cr. These results confirm the suitability of the identified areas for the collection of *Cystoseira* spp. thalli to be subjected to laboratory Cr enrichment, to exclude the effects on the spectral and photosynthetic response due to a natural photophysiological adaptation to high levels of Cr.

**Fig. 4.**
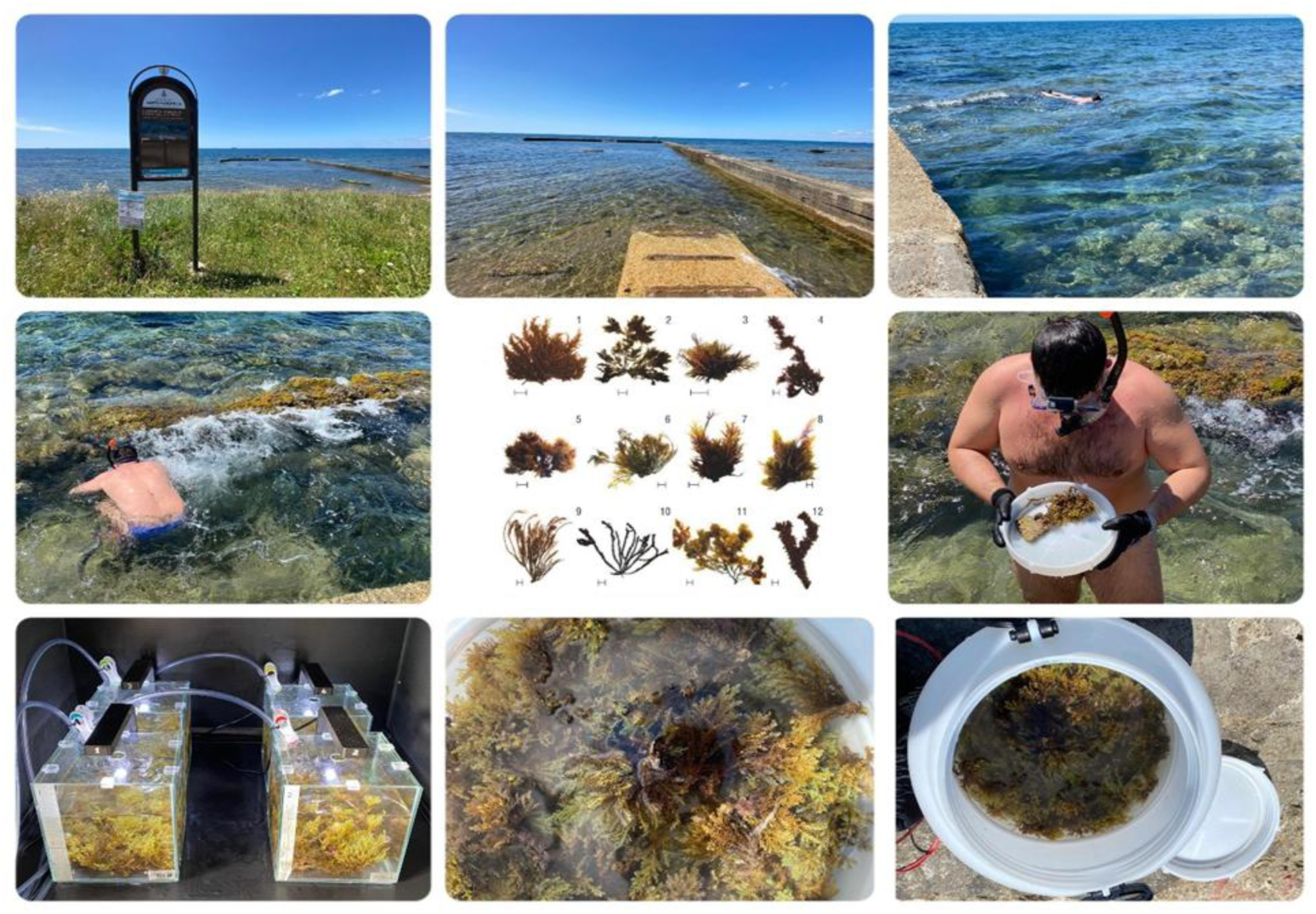
Sampling site of *Cystoseira* spp.’Molo L’ Roman structure, Civitavecchia SUD (42.05034 N, 11.81955 E)

Area 4

This is an area about 7.5 km south of the port. As in the previous area, monitoring was conducted along transects parallel to the coast. Here too, the presence of *Cystoseira* spp. was not observed in the stretch of the south coast of the site’Le Due Palafitte’.

## 5. Sampling site “Area 3”

Thalli used for the experiment was collected at 20-50 cm depths from the coastal area indicated in figure 4, in which the presence of *Cystoseira* spp. was detected during previous campaigns. In addition, to exclude the effects on the spectral and photosynthetic response due to a natural photo-physiological adaptation to high Cr levels, n.3 replicates of *Cystoseira* spp. (500 g for each sample) and seawater samples (250 ml for each sample) were collected and immediately analysed for the detection of Cr concentrations according to the EPA 7199:1996 method for Total Cr and UNI EN ISO 15587-1 2002. The analysis confirms the suitability of the identified areas for the collection of *Cystoseira* spp. thalli to be subjected to Cr laboratory enrichment. Once at laboratory, thalli were acclimatized to the culture conditions in aerated seawater collected at the sampling site to ensure a standard starting point for experiments at 21-22 °C. The quantity of *Cystoseira* spp. to be taken for each tank is 200 g, considering the quantity of biological material needed for chemical analysis.

## 6. Materials and Methods

The experimental protocol was developed in accordance with the methodological references reported in (Malea et al., 2021; Falace et al., 2018; D3.7 Progetto ASI-STOPP; Douay et al., 2022).

The experimental design consisted of 4 tanks (4L): 1 control tank and 3 tanks containing 3 concentrations of Cr: 0.1, 1 and 10 μmol/L (Baumann et al., 2009) (fig.5). Cr concentrations obtained from potassium chromate K₂CrO₄ Puriss grade chemicals (Sigma-Aldrich), diluted with seawater.

**Fig. 5.**
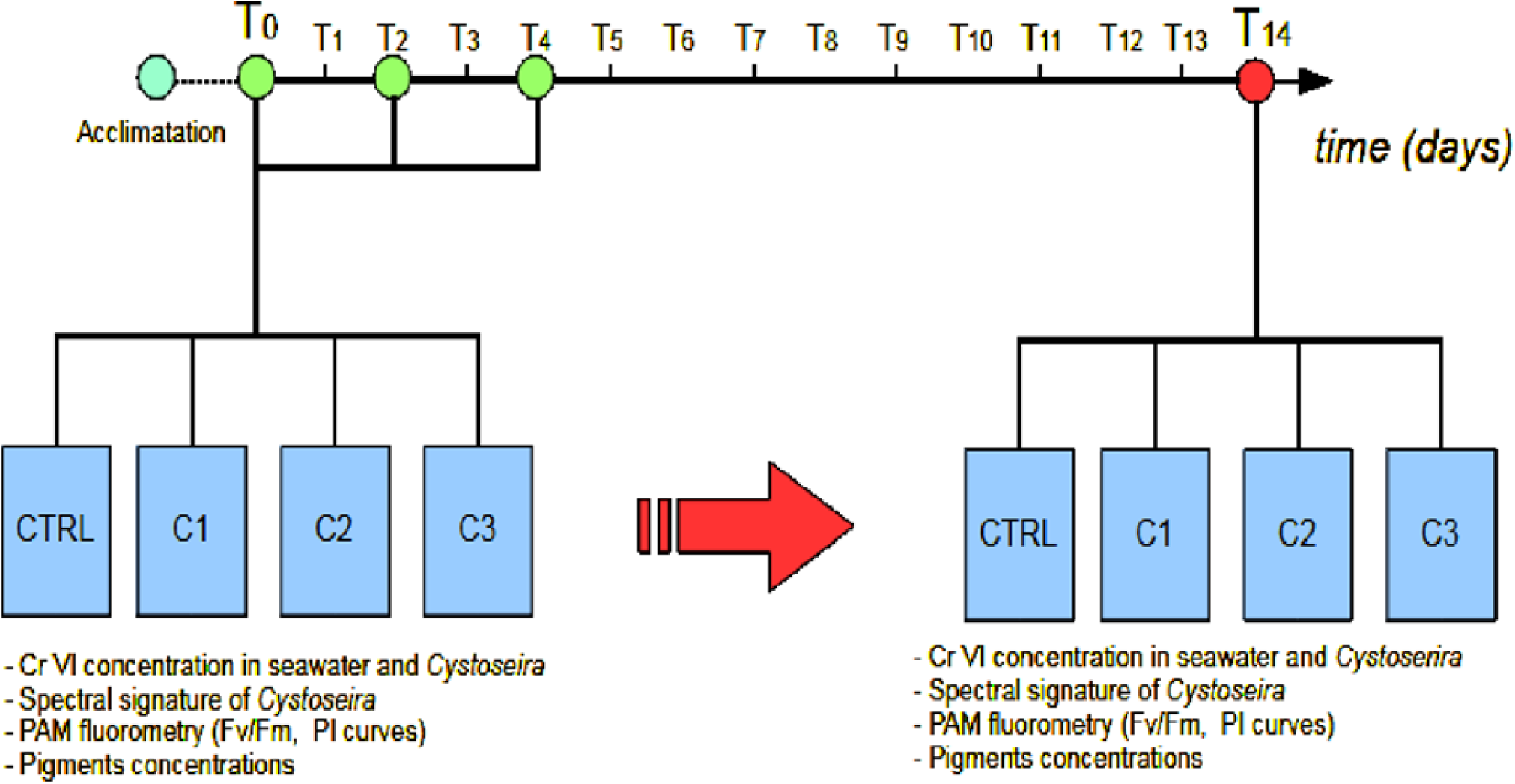
Experimental design protocol

Macroalgal cultures were maintained at a temperature range of 18-21 °C and a light regime of 120-125 μmol photon m⁻² s⁻¹. The experiment lasted 14 days, with measurements taken every two days during the first week and a final measurement at the end of the experiment (day 14).

At the beginning and end of the experiment, chromium analyses were performed on *Cystoseira* spp. and seawater by the analysis laboratory S.C.A. Servizi Chimici Ambientali s.r.l.

Each measurement cycle included the following types of measurement in 3 replicates for each tank (fig. 6):

**Fig. 6.**
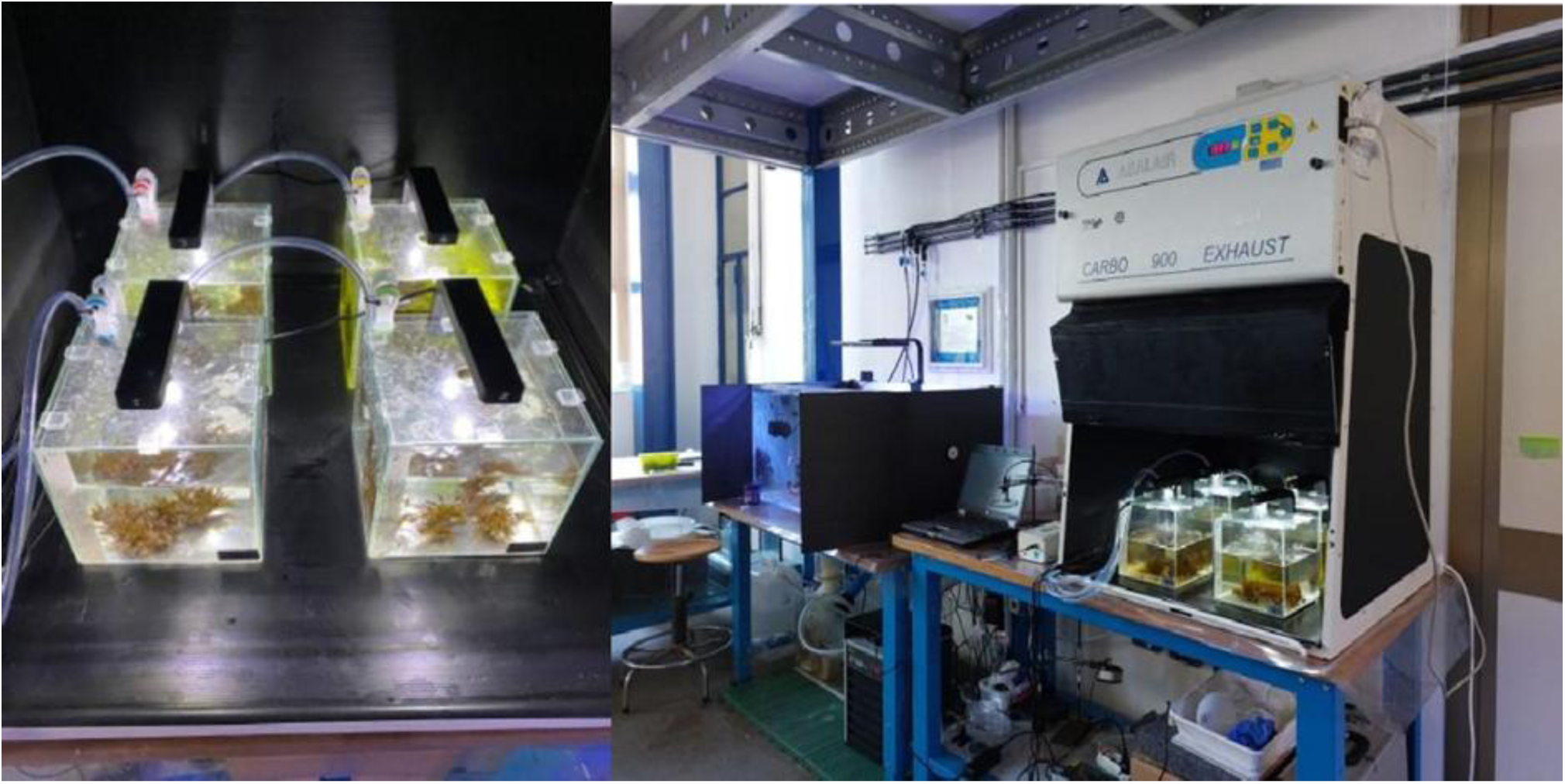
Experimental setup in the darkened hood with *Cystoseira* spp. Exposed to different concentrations of chromium

1. Spectrometric measurements were performed under clear sky conditions using the UV-VIS Stellarnet 2000 EPP spectrometer, equipped with fiber optics and a cosine corrector diffuser. The acquired parameters out of water were:

- water leaving spectral radiance (with the fiber optic orthogonally pointed to the surface of tank) obtained by using as incident light a halogen and a xenon lamp;

- reference incident radiance on a spectralon diffuser surface (with the same geometry used for the measurement of the water leaving radiance) of the light source;

- in air reflectance of *Cystoseira* spp. samples exposed to solar light during clear sky conditions (see Douay et al., 2022).

2. Fluorimetric and photosynthetic measurement by using the Junior PAM fluorometer (Waltz) according to Baumann et al., (2009) and Madonia et al., (2021):

A) Measurement of Fv/Fm (Maximum Quantum Yield of PSII)

‒ Dark adaptation: 15–20 minutes (using leaf clips or keeping samples in a dark chamber).
‒ Saturation pulse: ∼8,000 µmol photons m⁻² s⁻¹ for 0.8 seconds.
‒ Fluorescence recording:
• Fo (initial fluorescence) recorded using a weak measuring light (∼0.05 µmol photons m⁻² s⁻¹).
• Fm (maximum fluorescence) measured with the saturation pulse.
‒ 6. Calculation: Fv/Fm=(Fm−Fo)/Fm
B) Rapid Light Curves (RLCs)
‒ Number of light steps: 8–10 steps.
‒ Duration of each step: 10–20 seconds.
‒ Actinic light intensity intervals: typically between 0 and 1,000 µmol photons m⁻² s⁻¹, distributed logarithmically or linearly (e.g., 0, 10, 20, 50, 100, 200, 400, 600, 800, 1,000 µmol photons m⁻² s⁻¹).
‒ Saturation pulse: ∼8,000 µmol photons m⁻² s⁻¹ for 0.8 seconds at each step.
‒ Data recording:
• Y(II) = (Fm′−F)/Fm′, where F is the steady-state fluorescence and Fm’ is the maximum fluorescence under actinic light.
• ETR (Electron Transport Rate) can be calculated if the actinic light wavelength and absorption coefficient are known.

3. The spectrophotometric analysis of photosynthetic pigments was conducted following the methodology described by Stengel and Connan (2015). This method allows for the quantification of chlorophylls and carotenoids extracted from *Cystoseira* samples.

A) Sample Preparation and Pigment Extraction

1. Fresh algal samples were collected and immediately stored in darkness at-80°C until further processing to prevent pigment degradation.
2. Samples were homogenized in 90% acetone using a mortar and pestle under cold conditions to minimize pigment oxidation.
3. The homogenized samples were centrifuged at 4000 x g for 10 minutes at 4°C to separate the supernatant containing the extracted pigments.
4. The supernatant was carefully transferred to a clean tube, avoiding contamination with debris.

B) Spectrophotometric Measurements

The absorbance of the pigment extract was measured using a UV-Vis spectrophotometer.
Readings were taken at specific wavelengths corresponding to pigment absorption peaks:
‒ Chlorophyll a: 664 nm (corrected with 750 nm)
‒ Chlorophyll b: 647 nm (corrected with 750 nm)
‒ Chlorophyll c: 630 nm (corrected with 750 nm)
‒ Chlorophyll d: 696 nm (corrected with 750 nm)
‒ TotalChls = Chl-a+Chl-b+Chl-c+Chl-d

C) A blank measurement was performed using 90% acetone to calibrate the spectrophotometer.

Calculation of Pigment Concentrations:

The concentration of each pigment was calculated using the specific extinction coefficients provided by Stengel and Connan (2015):

‒ Chl-a = 11.85A664 – 1.54A630
‒ Chl-b = 20.13A647 – 5.03A664
‒ Chl-c = 24.52A630 – 2.99A6645
‒ Chl-d = 21.24A696 – 3.32A664
‒ TotalChls = Chl-a+Chl-b+Chl-c+Chl-d

where A represents the absorbance at the respective wavelengths. Quality Control and Replicates

- Each measurement was conducted in triplicate to ensure reproducibility.
- Pigment stability was monitored by minimizing exposure to light and heat throughout the process.
- Results were expressed as μg pigment per g fresh weight of algal tissue.

Data distribution was initially modelled by using the Shapiro-Wilk test (Shapiro and Wilk,1965); this preliminary test indicated that the data did not follow a normal distribution, therefore for further analyses a non-parametric model, the Kruskal-Wallis test (Kruskal and Wallis, 1952), was used. This latter test provides an effective statistical tool for comparing the features characterizing multiple samples and testing whether they fit with the same distribution. Furthermore, Wilcoxon-Mann-Whitney test (Fagerland and Sandvik, 2009) was used as two-sample test.

## 7. Results

The spectra reported in Figure 7 effectively show the optical responses of *Cystoseira* spp. under different experimental conditions at different Cr concentrations. In the left column, the irradiance curves are presented for four experimental conditions: the red line (CTRL) corresponds to the control condition with 0 μmol Cr, the blue line (C1) represents

**Fig. 7.**
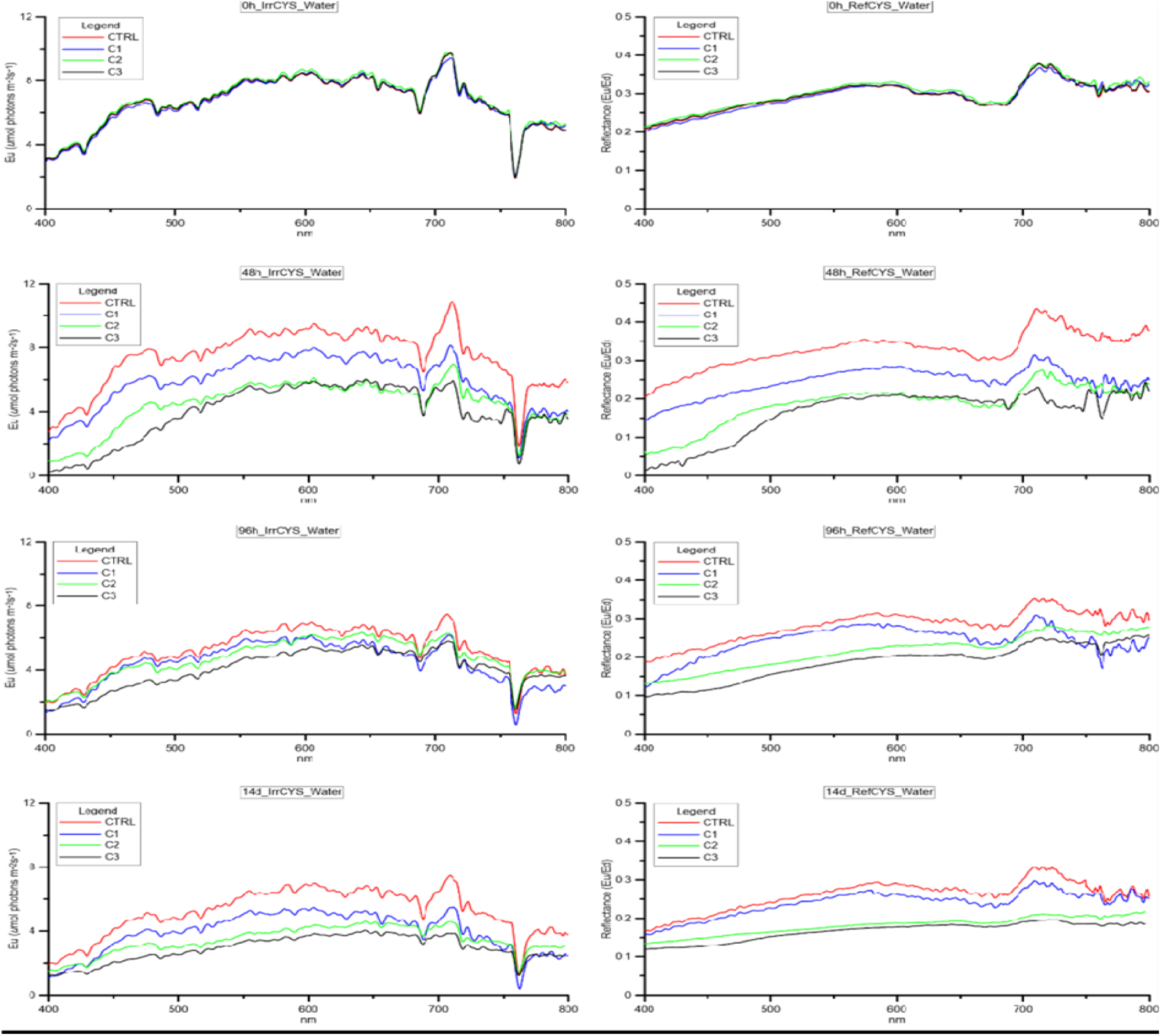
The figure is divided into two columns: the left column represents the irradiance measurements, while the right column illustrates the reflectance curves of *Cystoseira* spp. during the experiments with Chromium.

0.1 μmol Cr, the green line (C2) corresponds to 1 μmol Cr, and the black line (C3) represents 10 μmol Cr. Each plot captures the optical response of *Cystoseira* spp. at specific time points of the experiment: 0 h, 48 h, 96 h, and 14 days. The y-axis of these plots is expressed in units of photon flux density (µmol photons m⁻² s⁻¹), while the x-axis expresses the wavelength in nm.

In the right column, reflectance curves are shown for the same experimental conditions and time steps. The R(λ) on the y-axis describes how the incident light interacts with the algal surface, providing valuable information on the structural and pigment changes that occur over time.

The spectra are characterized by two key features that are noteworthy in the context of benthic vegetation and optical properties of brown algae. First, there is a prominent peak at 570 nm, which is indicative of accessory pigments such as fucoxanthin, a pigment unique to brown algae. Fucoxanthin plays a critical role in photosynthetic efficiency by absorbing light in the blue-green range, ensuring optimal energy capture under specific environmental conditions.

Second, the spectra reveal a pronounced peak at 710 nm, which corresponds to the red-edge effect, a well-documented phenomenon in algal studies. This effect marks the sharp transition between the strong absorption of chlorophyll in the visible red region (about 680 nm) and the high reflectance in the near-infrared (NIR) region (∼710 nm). In benthic vegetation such as *Cystoseira* spp., this feature reflects the internal structure of the algae and its physiological responses to stress or environmental changes, including chromium exposure.

The graph (Fig.8) represents the normalized spectral reflectance response of a *Cystoseira* spp. sample over a wavelength range from 400 to 800 nm after 14 days of exposure to chromium.

**Fig. 8.**
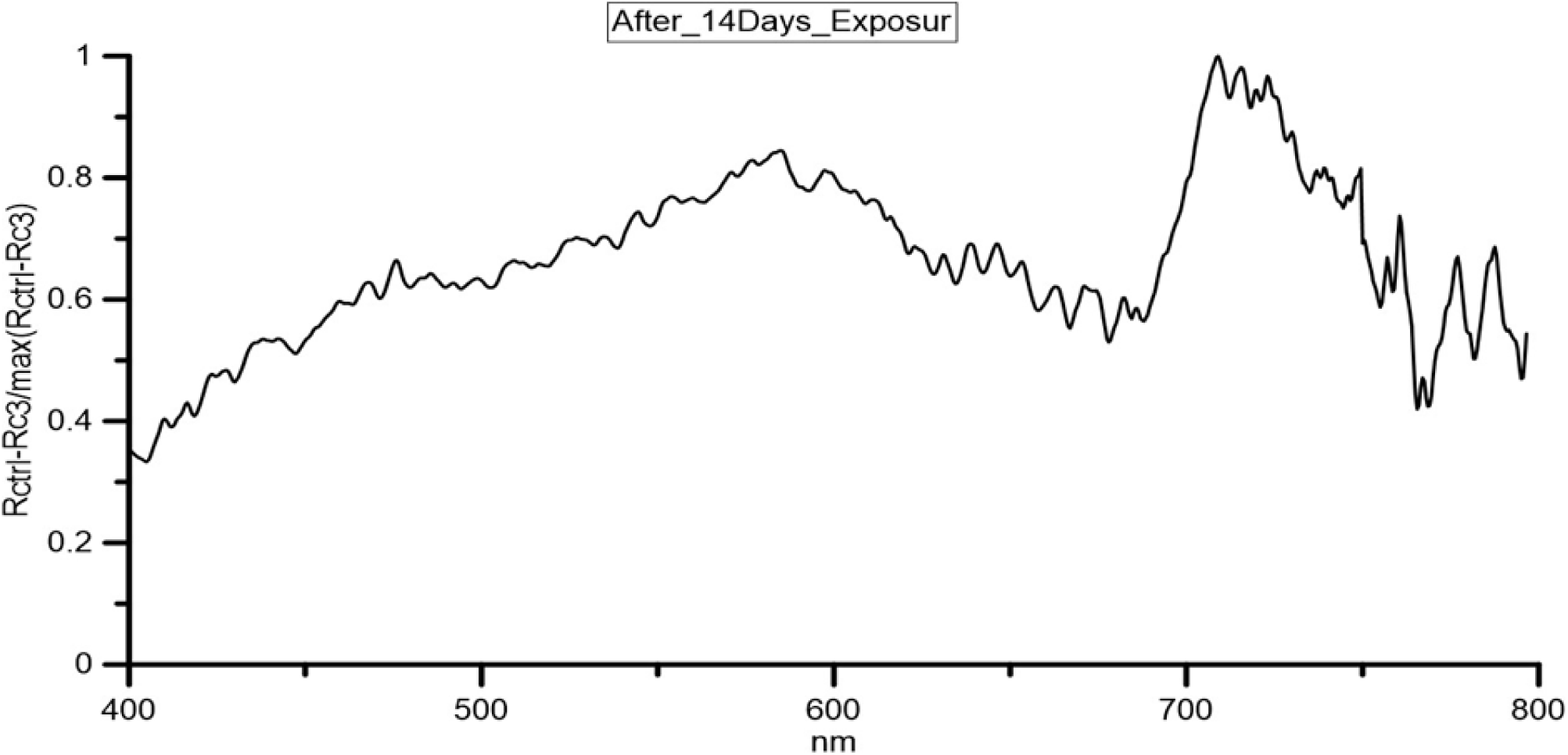
*Cystoseira* spp. Rctrl-Rc3/max (Rctrl-Rc3) after 14 days of exposure to chromium.

The y-axis indicates the normalized reflectance value, where the difference in reflectance between the control (Rctrl) and the sample with the highest chromium concentration (RC3) has been divided by the maximum value of this difference, ensuring that the data are dimensionless and that the values are between 0 and 1. The x-axis corresponds to the wavelength of light in nanometers (nm), which extends from the visible spectrum to the near-infrared region.

The graph shows characteristic spectral features, such as a large increase in reflectance between 500 and 600 nm, followed by a sharp peak near 700 nm. These features correspond to specific absorption or reflectance characteristics of the pigments or chlorophyll fluorescence in the sample. These reflectance data contain small oscillations, which indicate potential noise or natural variability in the optical properties of the sample. By normalizing the reflectance, the plot highlights relative spectral differences without being influenced by absolute quantities, making it suitable for comparative analysis.

After 14 days, the difference between the control and the C3 tank, with the highest Cr concentration, reached the maximum value, above all for the spectral region higher than 710nm that confirming the usefulness of this wavelengths also for marine mesolittoral vegetation monitoring.

The photosynthetic efficiency (P-E) curves presented in Fig.9 provide a detailed assessment of the light-dependent photosynthetic performance of *Cystoseira* spp. under different experimental conditions. These curves plot the relationship between photosynthetically active radiation (PAR, µmol photons m⁻² s⁻¹) on the x-axis, representing the range of light wavelengths available for photosynthesis (400– 700 nm), and electron transport rate (ETR, µmol e⁻ m⁻² s⁻¹) on the y-axis, an indicator of the photosynthetic electron flow through photosystem II (PSII).

**Fig. 9.**
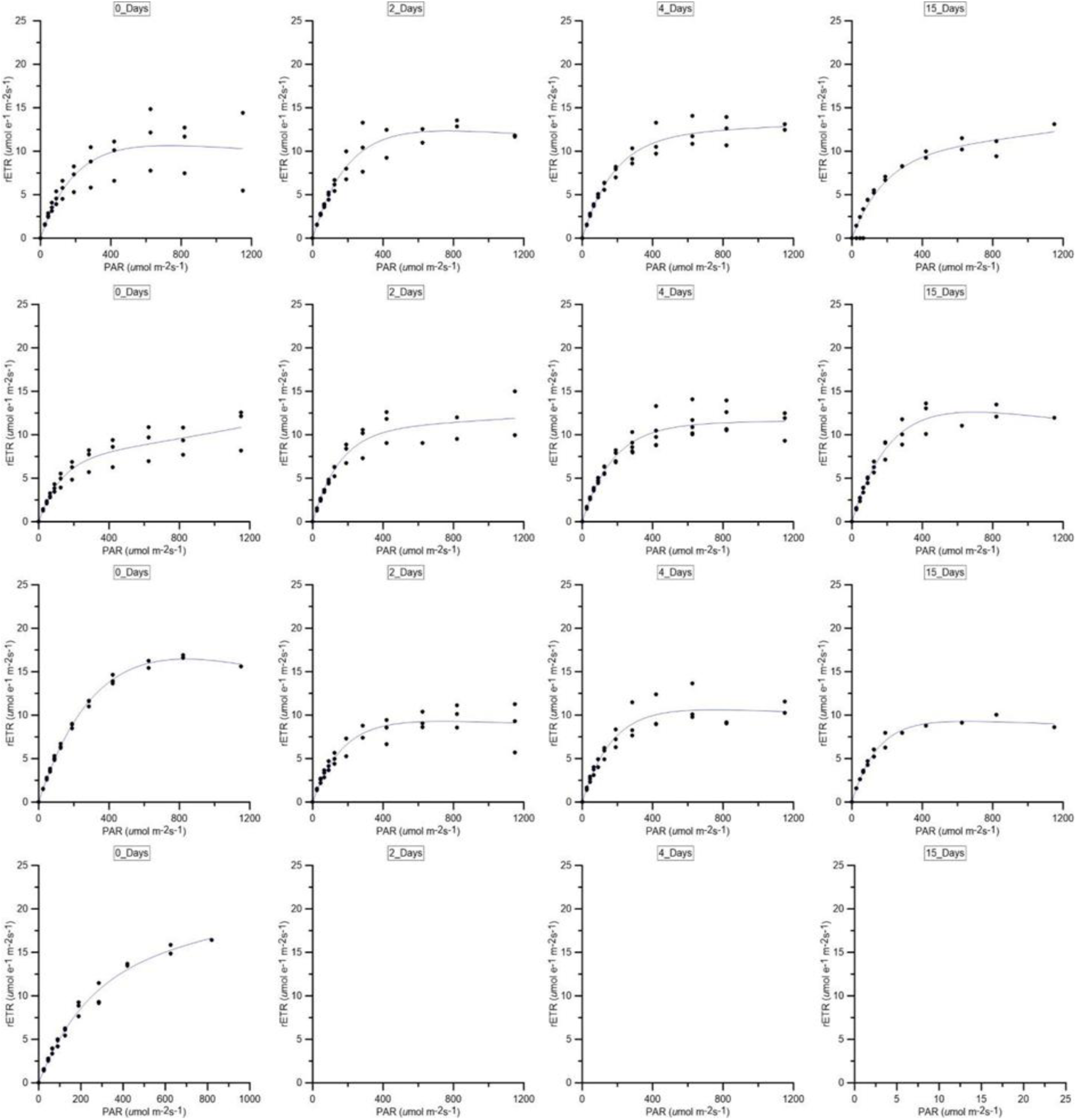
Cystoseira spp. RLCS before and after the exposure to Chromium (PAM measurements).

To verify the influence of Cr concentrations on the algae over time, and the presence of statistically significant differences among the data, statistical tests (Gower’s distance PERMANOVA) were conducted considering the Cr concentrations in the tanks water and in the algae thalli, the bioaccumulation factor (BAF), the concentrations of photosynthetic pigments (chla, chlb, chlc, and chld) in the *Cystoseira* spp. thalli, the variable fluorescence/maximum fluorescence in shaded (Fv/Fm) leaves indicates variations in the photonic efficiency of photosystem II (PSII) (Adams et al., 1995) (Table 1) and the Electron Transport Rate (ETR) (Fani and Lazzara, 2006) (Fig.9).

**Table 1.** Cystoseira spp. spectral signature before and after the exposure to Chromium.

| ID | chl a (µg/ml) | chl b (µg/ml) | chl c (µg/ml) | chl d (µg/ml) | Totals Chls (µg/ml) | Fv/Fm |
| --- | --- | --- | --- | --- | --- | --- |
| CTRL_0 | 8,217 | 0,054 | 1,055 | 0,223 | 9,437 | 0,685 |
| CTRL_2days | 9,673 | 0,168 | 1,391 | 0,228 | 11,327 | 0,655 |
| CTRL_4days | 12,980 | 0,206 | 1,565 | 0,249 | 14,820 | 0,698 |
| CTRL_14days | 8,731 | 0,268 | 1,416 | 0,340 | 10,635 | 0,644 |
| C1_0 | 10,587 | 0,152 | 1,488 | 0,235 | 12,312 | 0,644 |
| C1_2days | 11,795 | 0,272 | 1,548 | 0,255 | 13,706 | 0,659 |
| C1_4days | 10,974 | 0,201 | 1,469 | 0,218 | 12,711 | 0,684 |
| C1_14days | 14,019 | 0,190 | 1,788 | 0,234 | 16,040 | 0,658 |
| C2_0 | 13,414 | 0,336 | 1,780 | 0,283 | 15,629 | 0,631 |
| C2_2days | 10,589 | 0,048 | 1,397 | 0,221 | 12,111 | 0,649 |
| C2_4days | 9,031 | 0,120 | 1,098 | 0,161 | 10,287 | 0,635 |
| C2_14days | 5,616 | 0,085 | 0,801 | 0,125 | 6,548 | 0,571 |
| C3_0 | 8,752 | 0,135 | 1,172 | 0,174 | 10,111 | 0,645 |
| C3_2days | 6,453 | 0,487 | 0,854 | 0,224 | 7,929 | 0,000 |
| C3_4days | 4,435 | 0,454 | 0,410 | 0,190 | 5,430 | 0,000 |
| C3_14days | 1,547 | 0,424 | 0,163 | 0,137 | 2,250 | 0,000 |

In general, the experiments clearly showed that as the Cr concentrations in the water increased (tanks CTRL>C1>C2>C3), the bioaccumulation factor (BAF) decreased (C3<C2<C1<CTRL).

Considering the photosynthetic pigments (chla, chlb, chlc and chld), a statistically significant difference in long-term chlc concentrations emerged (KW test; pvalue < 0.05) considering all the tanks. A statistically significant difference emerged also considering the Fv/Fm ratio values between the four tanks in the long term (KW test, p value < 0.05) and this difference was particularly marked between the values measured in CTRL and C3 (MW test; p value <0.05).

To evaluate the interaction between all the variables measured during the experiment, the multivariate analysis PERMANOVA was applied (Anderson, 2014). In the PERMANOVA analysis, the Gower distance and 9999 permutations were used.

This distance was chosen because it allows for quantifying the dissimilarity between samples characterized by multiple types of data (e.g., categorical, ordinal, and continuous traits; Mouillot et al., 2021), eliminating the need for preliminary transformation to a common scale (De Bello et al., 2021). The results showed a statistically significant difference between CTRL and C3 (p < 0.05), considering Cr concentrations in water, BAF, photosynthetic pigment concentration, Fv/Fm ratio values, and median ETR values.

## 8. Discussion and Conclusions

The present study provides novel insights into the reflectance and irradiance properties of *Cystoseira* spp. across different experimental conditions. Our findings align with recent work on macroalgae spectral characteristics, highlighting the importance of environmental factors in influencing light absorption and reflectance patterns (Smith et al., 2015; Roberts et al., 2017). The spectral behavior observed in our study closely resembles patterns identified in related species, such as Sargassum and Fucus, which also exhibit adaptive responses to varying light regimes (Polo et al., 2015).

One of the most striking results is the significant variation in reflectance at key wavelengths (e.g., 450– 550 nm and 650–700 nm), which suggests a dynamic photophysiological response to environmental stressors. These findings are supported by the work of Lee, Z. et al. (2020), who demonstrated that changes in reflectance spectra are strongly correlated with pigment composition and photoacclimation strategies in brown algae. Furthermore, spectral shifts at longer wavelengths (>700 nm) could indicate alterations in structural properties, such as changes in cell wall composition and surface roughness (Álvarez-Gómez et al., 2017). These structural changes have been extensively documented in studies on macroalgal responses to hydrodynamic forces and nutrient availability (Spalding et al., 2019).

These observations are consistent with the findings in the literature, in fact according to Nourisson et al. (2016), the red-edge effect is a diagnostic feature in benthic vegetation, providing information on pigment composition and structural adaptations. Furthermore, Michaelian (2015) highlights how reflectance changes in algae are influenced by structural and physiological processes, evident in the spectra presented here. Finally, data are available on the bioaccumulation of some trace metals in the soft tissues of *Paracentrotus lividus* (Lamarck, 1816) in the study area, which basically confirm the influence of sediment metal concentrations on the distribution of bioaccumulation levels in the sea urchin (Scanu et al., 2015).

The differences observed between in-air and in-water measurements confirm the significance of immersion effects on spectral reflectance. Our results corroborate previous research by Souri (2019), who showed that water column properties, including turbidity and dissolved organic matter, can influence spectral signatures in submerged macroalgae. Additionally, optical modeling studies emphasize the role of water depth and dissolved particulate matter in modifying reflectance profiles, which is consistent with our observations (Hamidifar et al., 2024). This has crucial implications for remote sensing applications, as highlighted by Brown et al. (2022), who emphasized the necessity of accounting for environmental interference in spectral measurements of aquatic vegetation.

An interesting study is the work of Hoang et al. (2016), who produced a submerged aquatic vegetation (SAV) spectral signature library to collect optical data of macroalgae (red, green and brown), seagrasses and sediment characteristics in coastal waters, to support the selection of spectral bands and bandwidths for different environmental conditions, such as clear water and high turbidity water bodies, and to choose satellite sensors suitable for different benthos. Twenty-two species of submerged coastal aquatic plants, including red, green, brown macroalgae and seagrasses, were collected at Cape Peron, Western Australia, to measure their spectral reflectance in situ.

Moreover, our findings provide additional evidence of the role of photoprotection mechanisms in brown algae. The observed increase in reflectance within the 500–600 nm range under stress conditions aligns with the work of Seys et al. (2022), who found that xanthophyll cycle activation and increased carotenoid content contribute to the modulation of reflectance under high irradiance levels. Such mechanisms are vital for mitigating oxidative stress and ensuring algal survival in fluctuating light environments. Additionally, studies by Melo-Santos et al. (2018) have shown that brown algae exposed to environmental stressors such as salinity changes and UV radiation develop photoprotective adaptations that are reflected in their spectral properties.

Our study also highlights the potential use of spectral signatures for detecting physiological stress in *Cystoseira* spp. Under conditions of prolonged stress, the decline in near-infrared reflectance could be indicative of cellular damage and chlorophyll degradation, a phenomenon previously reported by Zahir et al., (2022) in seagrasses and macroalgae. This suggests that hyperspectral reflectance analysis could serve as an early-warning system for assessing macroalgal health in response to environmental disturbances, a crucial tool for conservation strategies Wahl et al., 2014).

In conclusion, our study demonstrates the intricate relationship between spectral reflectance properties and environmental conditions in *Cystoseira* spp. The observed spectral variations, both in-air and in-water, highlight the photophysiological adaptability of these macroalgae, in agreement with recent literature (Polo et al., 2015; Lee et al., 2020; Seys et al., 2022). These findings have significant implications for the development of remote sensing methodologies aimed at monitoring macroalgal health and productivity.

Future research should focus on refining spectral models to improve the accuracy of remote sensing applications, particularly in distinguishing stress-induced spectral signatures from natural variability (Álvarez-Gómez et al., 2017). Additionally, integrating physiological and biochemical analyses with spectral data could provide deeper insights into the adaptive strategies of *Cystoseira* spp. under different environmental stressors (Brown et al., 2022). The inclusion of long-term monitoring approaches, such as those proposed by Carvalho (2019), could further enhance our understanding of macroalgal responses to climate-driven environmental changes.

Moreover, advances in hyperspectral imaging and machine learning algorithms offer new opportunities for the automated classification of spectral data, improving the detection and quantification of algal biomass in coastal ecosystems (Zhang et al., 2022). Such advancements could revolutionise ecological assessments, aiding in habitat conservation and marine resource management.

Ultimately, this study contributes to the growing body of research on macroalgal spectral properties, reinforcing the importance of spectral monitoring in ecological assessments and conservation efforts (Souri, 2019). By advancing our understanding of algal spectral responses, we can enhance the effectiveness of marine ecosystem management strategies in the face of global environmental changes. The continued development of spectral monitoring techniques will be instrumental in ensuring the resilience of marine macroalgal communities in the coming decades (Wahl et al., 2014).

